# Development and evaluation of a low-cost, automated camera trap for surveying bumble bee communities

**DOI:** 10.64898/2025.12.09.692866

**Authors:** Michael P. Getz, Lincoln R. Best, Oksana Ostroverkhova, Andony P. Melathopoulos, Timothy L. Warren

## Abstract

Widespread declines in insect diversity and abundance underscore an urgent need for standardized, nonlethal monitoring methods for important pollinators such as bumble bees (*Bombus* spp.). Camera traps are widely used for non-intrusive, continuous surveys of large animals but have not yet been extensively adopted for monitoring insects. An automated camera trap system could improve current insect survey methods, which often rely on lethal traps or in-person observation. We developed an open-source, low-cost camera trap for monitoring wild insects and evaluated its performance relative to established sampling approaches. The system used a low-power microcomputer to collect time-lapse images on colored platforms. We trained whole-image and tiled deep-learning object detection models to detect insects in images captured by the traps. We found that tiled inference models significantly improved detection accuracy and outperformed human review. Bumble bee visitation was highest on platforms featuring a fluorescent bullseye pattern; adding a fluorescent coat to blue platforms increased visitation modestly. With continuous monitoring, the cameras recorded bumble bee visits during all daylight hours. Over 18 days, camera traps recorded six *Bombus* species, yielding community composition and diversity estimates comparable to those obtained by hand netting and blue vane traps. Using our observed data, we simulated the effect of deploying variable numbers of cameras at sites with distinct levels of diversity. Adding cameras substantially increased sampling completeness, particularly in species-rich communities. Our findings demonstrate that low-cost, automated camera traps paired with deep-learning image analysis can enable scalable, nonlethal studies of bumble bee diversity and behavior. Our work establishes a foundation for monitoring other diurnal insect communities.

## Introduction

There is considerable evidence of recent, worldwide declines in the diversity and abundance of insects (Sánchez-Bayo and Wyckhuys, 2019; Van Der Sluijs, 2020; Wagner, 2020). Of particular concern are North American and European native bees, especially bumble bees (*Bombus* spp.), which are effective and economically important diurnal pollinators (Biesmeijer et al., 2006; Cameron et al., 2011; Cameron and Sadd, 2020; Jacobson et al., 2018). For many native bees, however, trends in range and abundance are not well understood; even at a coarse scale of 220x220 km, reliable inventory data exists for fewer than 40% of bee species in the United States (Chesshire et al., 2023). These gaps reflect limitations of conventional monitoring techniques (e.g. aerial netting, temporary immobilization), which can be biased, poorly suited for detecting rare species or community changes, and involve highly invasive specimen handling (Eeraerts and Meeus, 2025; McNeil et al., 2019; Montero-Castaño et al., 2022; Williams et al., 2001).

Passive, lethal trapping methods are largely prohibited for species of concern and rely on overleveraged bee taxonomists to process specimens (Mathis et al., 2024; Portman et al., 2020; Portman and Tepedino, 2021). Therefore, there is an urgent need to develop automated, nonlethal approaches to bumble bee monitoring.

Camera-based systems show potential for standardized, nonlethal insect monitoring. In larger animals, cameras often trigger image capture with passive infrared sensors, which are less reliable for small, ectothermic insects (Boom et al., 2014; Rovero et al., 2013; Rowcliffe et al., 2014; Samejima et al., 2012). Continuous, scheduled imaging is suitable for insects but generates large volumes of data without specimens (Droissart et al., 2021; Naqvi et al., 2022). To address this problem, recent studies have developed deep learning detection models to localize insects, often using simplified backgrounds, such as immobile, low-lying plants (Bjerge et al., 2022) or stationary, attractive lures (Sittinger et al., 2024; Teixeira et al., 2023). Object detection models have achieved high accuracy across multiple insect groups (Bjerge et al., 2023; Sittinger et al., 2024), but may not generalize to different camera platforms or taxa. Specialized detection methods for small objects, such as tiling, may aid insect detection, but it is unclear whether performance gains merit the added computational effort required (Ye et al., 2024). Bumble bees are ideal targets for image-based monitoring given their relatively large size, recognizable markings, and ecological significance.

To effectively survey an insect community, a passive trap must attract as many species as possible. Light traps are highly effective for nocturnal insects such as moths (Montgomery et al., 2021), but attracting diurnal insects, which rely on color, is more challenging. Certain colors and patterns are known to attract bumble bees. For example, lethal blue vane traps are highly attractive to *Bombus* spp. (Hall, 2018; Rao and Ostroverkhova, 2015; Stephen and Rao, 2005), owing in part to the traps’ fluorescence and reflectance at 430 nm, aligned with the peak sensitivity of a bee photoreceptor (Ostroverkhova et al., 2018). Many plants attract bumble bees via UV-reflective signals, including a concentric bullseye pattern, indicating pollen availability or potential floral rewards (Gronquist et al., 2001; Guldberg and Atsatt, 1975; Koski and Ashman, 2014). The contrast between floral patterns and the background may further allow bees to identify targets (Hempel De Ibarra et al., 2022; Van Der Kooi et al., 2019). The observed efficacy of blue vane traps and high-contrast floral guides suggests that combining these cues in a blue-fluorescent bullseye pattern on a platform lure could attract bumble bees.

The extent to which automated imaging systems can survey bee communities is not well understood. There are substantial differences in bee community estimates obtained by blue vane traps, pan traps, and hand netting. Netting and vane traps tend to overrepresent *Bombus* spp., whereas pan traps more often collect small halictids (Boyer et al., 2020; Mathis et al., 2024; Wilson et al., 2008). Vane traps of different colors attract distinct *Bombus* species (Acharya et al., 2021). Such discrepancies highlight the need to validate community estimates obtained by camera traps, using methods that account for unequal sampling effort (Chao et al., 2020; Westphal et al., 2008).

Here we developed an open-source, time-lapse camera trap system for imaging insects on static backgrounds. We compared two custom-built deep learning insect detection models and found that tiling images into smaller, 640 x 640 pixel regions significantly improved bumble bee detection. We observed that bumble bees visited the fluorescent bullseye platform more than platforms with solid colors that constitute the bullseye pattern. Our camera systems estimated the diversity of bumble bee communities as effectively as hand netting, without destructive sampling. Finally, using a simulation approach, we compared sampling efficiency with variable numbers of camera traps at low and high diversity sites.

## Materials and Methods

### Camera traps for pollinator monitoring

We constructed ten camera traps using readily available materials at a per-unit cost of about US$190. The traps were controlled by low-power (∼1-1.5 W) Raspberry Pi Zero or Zero 2W computers (Raspberry Pi Foundation, Cambridge, UK), which ran Bullseye 64-bit Lite from a 64 GB Secure Digital (SD) card (SanDisk, Milpitas, CA, USA). A 19,200 mAh Voltaic Systems V75 battery in a dry bag provided power, and a Witty Pi 4 Mini controlled startup and shutdown (Voltaic Systems, Brooklyn, NY, USA; Dun Cat B.V., Elst, NL). An OLED display (Zhong JinYuan Technology, Henan, CN) and precision real-time clock (Adafruit, Brooklyn, NY) connected to the Pi via a SparkFun Qwiik Hat (SparkFun, Niwot, CO). The electronics were housed in a weatherproof box mounted to an aluminum frame shaped like an inverted L. A Pi Camera Module 3 (Raspberry Pi Foundation, Cambridge, UK) in a protective case was mounted to the top arm of the frame and pointed down at the platform lure. The frame was clamped to a 61 cm rebar stake hammered into the ground.

We replaced SD cards and batteries roughly every 48 hours. Imaging occasionally paused due to hardware malfunctions, adverse weather, or delayed site visits. All traps initially used the Pi Zero computer, imaging at 0.5 Hz. Two traps were upgraded to the Pi Zero 2W on day 3, which imaged at 1 Hz. On day 6, Pi Zero 2Ws were installed on all traps, standardizing imaging to 1 Hz. On day 6, we also extended the imaging hours from 5:00 AM - 4:00 PM to 6:00 AM - 8:00 PM to increase daylight coverage. Imaging at 1 Hz for 14 hr/day, data storage and battery power lasted ∼72 hours. Images were captured at 2304 x 1296 pixels with the focal plane set just above the platform. Camera trap software is available at https://github.com/warren-lab/bee_cam_basic.

### Platform variants

We developed five platform variants: green, yellow, blue, fluorescent blue, and a bullseye pattern. Platforms were 35.6 cm x 20.3 cm rectangles of white corrugated plastic, printed with color ink. To make a blue-fluorescent clear coat, we dissolved Risk Reactor Clear Blue Fluorescent Solvent Based Dye (Risk Reactor, Santa Ana, CA, USA) in methyl ethyl ketone (MEK), mixing the solution into clear nail polish (Sally Hansen Xtreme Wear, Invisible; Denton, TX, USA). The two fluorescent platforms were blue platforms coated with this solution. Green, yellow, and blue platforms corresponded to colors used in prior studies (Cane et al., 2000; Ostroverkhova, 2018; Acharya, et al., 2021). The bullseye platform had a green background with an array of blue rings (2.7 cm diameter) painted with fluorescent clear coat around yellow circles (1.5 cm diameter). The constituent colors in the bullseye platform matched those in single-color platforms. We used a repeating pattern because patches of colored circles are more detectable by bees than solitary disks (Wertlen et al., 2008). We reapplied the clear coat to the fluorescent blue and bullseye platforms twice in 18 days.

### Spectroscopy

We measured reflectance of the platform variants in the 350-1000 nm range with an Ocean Optics Spectrometer using a white lamp with tungsten filament (Ocean Optics, Orlando, FL, USA). Sample reflectance was normalized by subtracting the background. We measured photoluminescence (PL) of our fluorescent clear coat using an Ocean Optics USB2000-FLG Spectrometer with 405 nm laser excitation and a 410 nm long-pass filter. We averaged PL spectra from three locations on each sample and corrected for detector sensitivity using the AM1.5 Direct Solar reference (PV Education). Nail polish alone showed no PL under 405 nm excitation. We measured PL decay of the fluorescent clear coat by positioning a sample platform 2.5 cm under a UV lamp with emission recorded every 1-2 days, then weekly until stable. We estimated field-equivalent exposure by comparing the integrated UV energy (300-400 nm) of the AM1.5 solar spectrum with the UV lamp, yielding a conversion ratio from lamp time to field-exposure days. Normalized fluorescence intensity declined to 31.7% of its initial value after 8 exposure days and to 13.4% after ∼57 days. The decay was modeled as PL(t) = ae^-kt^ + c, where t is exposure time in days. Fitting this model to the normalized data yielded a decay constant of k = 0.254 day^-1^. We compared blue platform reflectance and clear-coat fluorescence to published blue vane trap spectra by independently normalizing the 400-600 nm ranges and assessing relative peak positions and spectral shape (Ostroverkhova et al., 2018).

### Sampling protocol

We deployed the ten camera traps in a red clover (*Trifolium pratense*) seed field at Hyslop Field Research Laboratory (Oregon State University, Corvallis, OR) during peak bloom (11-29 July, 2024). Red clover is highly attractive to bumble bees, and clover fields at this location and sites nearby have been the focus of prior bumble bee monitoring studies (Anderson et al., 2010; Rao & Stephen, 2009; Rao & Stephen, 2010). Platforms were randomly assigned and re-randomized on 18 and 23 July. From 25 July to 2 August, we deployed blue vane traps along the field edge ∼200 m east of the cameras. The blue vane traps had clear jars without liquid and were emptied every ∼48 hours; specimens were frozen, pinned, and identified. We conducted hand-netting on 15 July with one observer and on 1 August with four observers. Following established protocol (Krahner et al., 2021), samplers walked transects in clover plots for 10 minutes, collecting any bees within 1 m width. Bees were euthanized with ethyl acetate, pinned, and identified by OSU expert taxonomists.

### Image processing

We trained whole-image and tiled insect detection models on a computer (Ubuntu 22.04) with a single A4500 GPU (Nvidia, Santa Clara, CA, USA). To reduce false positives, we fine-tuned the pre-trained YOLO11s network (Jocher et al., 2023) using data from prior fieldwork and extensive background images. We reviewed detection output from the first five days of monitoring, adding undetected insects to the training dataset.

For the whole-image approach, images were scaled to 1280 x 1280 pixels (aspect maintained), which kept training batch size at 24 while preserving details of small insects. We split the training and validation set (72.3%/27.7%) using a date cutoff. Augmentations included flipping, scaling, and mosaic, which generates composites from the corners of four random images. We ran this initial model on the full camera trap dataset and reviewed detections to find images containing bumble bees. We trained a final model that included newly detected bumble bee images (70%/30% split). We duplicated early false positives (background areas misclassified as insects) and false negatives (missed insects) to improve discrimination. The final training set included 3,103 annotated images and 3,232 backgrounds; the validation set contained 296 bumble bee-only images.

To create the tiling dataset, we split full images into 640 x 640 tiles with 20% overlap. Bounding boxes were preserved across tile borders; tiles with small bounding box fragments were removed. We pruned redundant background tiles using image hashing. The tiled training set contained 6,162 images with full or partial insects and 57,095 empty tiles; the validation set had 633 bumble bee tiles and 2,436 empty tiles. Training parameters other than image size were identical to whole-image training.

We applied the trained models to all camera trap images on the Oregon State High Performance Computing cluster using a node with eight Nvidia GeForce GTX 1080 GPUs (Nvidia, Santa Clara, CA, USA). For the tiling model, we used Slicing Aided Hyper-Inference (SAHI; Akyon et al., 2022) to perform detection in a 640 x 640 px window moving stepwise across each full-resolution image, with overlap to ensure full coverage. Adjacent tile predictions were merged by combining conjoined bounding boxes; redundant detections were removed with non-maximum suppression. SAHI deductions were reported relative to the original, full-scale image.

We evaluated performance on the bumble bee-only validation set using recall and F1 score. Recall is the ratio of correct detections to all insects present. Precision is the ratio of correct detections to total detections. F1 score is the harmonic mean of recall and precision. We prioritized recall to minimize false negatives at the cost of including false positives, which were removed during manual review. We also compared model detections with those identified by a trained reviewer watching 30-frames-per-second videos of all images from a random day.

### Diversity Estimation and Sampling Completeness

We quantified species diversity and sampling completeness using Hill numbers and sample-coverage–based rarefaction and extrapolation, using the R package *iNEXT* (v3.0.2; Chao et al., 2014; Hsieh et al., 2016). Hill numbers unify classic diversity metrics by converting richness (q_0_, the count of species present), Shannon diversity (q_1_, which incorporates both richness and evenness of relative abundances), and Simpson diversity (q_2_, which emphasizes the dominance of common species) into “effective number of species,” placing these indices on a common, continuous scale (q ≥ 0). *iNEXT* generates standardized rarefaction and extrapolation curves, which estimate expected diversity at various sample sizes, with an asymptotic diversity estimate corresponding to infinite sampling. We assessed sampling completeness using coverage, the proportion of total community abundance represented by individuals sampled (Chao et al., 2014). Coverage-based rarefaction/extrapolation allows comparison across methods at equivalent completeness levels, regardless of differences in sample size (Chao et al., 2014).

### Modeling and Statistical Tests

We tested for differences in bee visitation among platform types by randomly permuting visit times among camera traps, counting visits only if the assigned camera was active at the time. We ran 10,000 iterations and adjusted P-values for multiple comparisons using the false discovery rate. We quantified the effect of fluorescence as the difference between simulated visit counts to the fluorescent-blue and blue platforms.

We modeled camera trap sampling efficiency for low- and high-diversity communities using two datasets: abundance data from this study (6 species), and Oregon Bee Atlas observations at a site near Bend, OR (13 species). For each site, we estimated mean sample coverage by bootstrapping species abundance data for 1, 2, and 3 cameras across 25 days, with 95% confidence intervals. We calculated coverage in Python using a function adapted from *PyNext* (https://github.com/davidlevybooth/pynext). We modeled observation rates with a negative binomial distribution fitted to visitation data from bullseye camera traps and randomly simulated hourly detections for a 14-hour sampling period.

### Software

We conducted analysis and visualization in Python (3.12.0) using packages *pandas* (2.1.1), *Matplotlib* (3.8.0), *Numpy* (1.26.1), *Seaborn* (0.13.0), *Scipy* (1.11.3), and *statsmodels* (0.14.5; Hunter, 2007; Seabold and Perktold, 2010; Virtanen et al., 2020; Waskom, 2021) Code is available at https://github.com/warren-lab/bee_cam_analysis.

## Results

### Image Collection

Over 18 days, our camera traps (Fig. 1A) collected a total of 5.7 million images (Fig. 1B). The platform types included three solid colors – blue, yellow, and green – and blue with a fluorescent clear coat (Fig. 1C). Additionally, a high-contrast platform had a green background and fluorescent blue rings around yellow centers, mimicking floral bullseye cues (Fig. 1C). We hypothesized that this high-contrast bullseye design would attract more bumble bee visits than solid colors. The peak reflectance of the blue platform was ∼430 nm, aligning with bee photoreceptor sensitivity (∼ 420-450 nm; Briscoe and Chittka, 2001; Peitsch et al., 1992) and blue vane trap reflectance (Fig. 1D). The blue vane trap reflectance, however, was much more narrowly distributed around the peak, indicating higher spectral purity. The green and yellow platforms both had peak reflectance at ∼520 nm, though the yellow reflected strongly at wavelengths up to 600 nm. The photoluminescence of the blue-fluorescent clear coat closely matched the blue vane trap, with a narrow peak at 430 nm (Fig. 1E).

**Figure 1:**
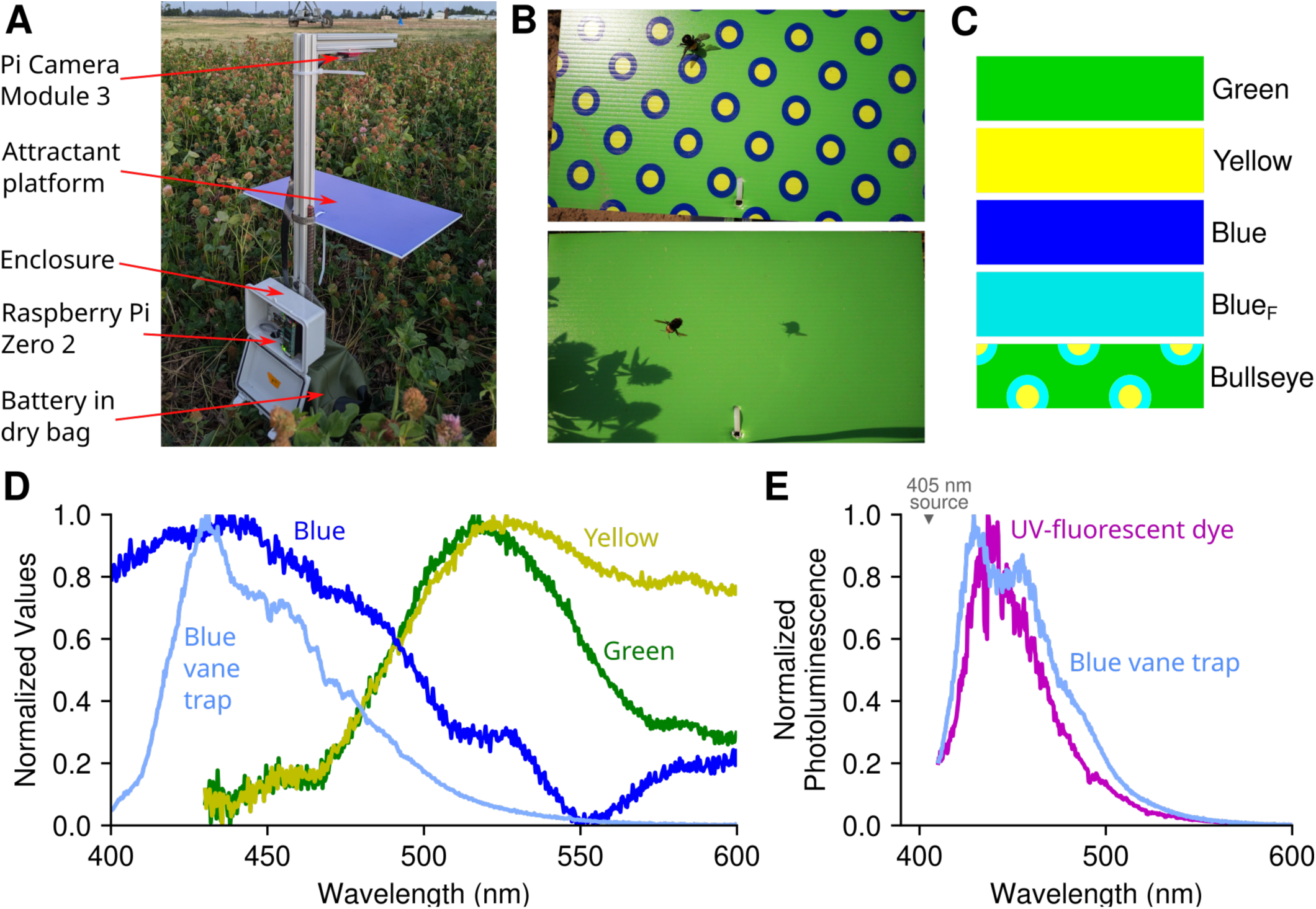
Design of an automated insect camera trap with non-rewarding attractant platforms. A) The custom-built camera trap was battery powered and controlled by a Raspberry Pi microcomputer. The camera was directed downward toward a platform with a printed color or pattern. B) Example images captured by a camera trap. C) We used five platform variants: blue, green, yellow, blue with fluorescent clear coat paint (Blue_F_), and a repeating bullseye pattern made up of green, yellow, and blue with fluorescence. D) Normalized reflectance spectra of the platform colors and a blue vane trap. The blue platform reflectance spectrum peaked near 430 nm, but was broader compared to the blue vane trap. The green platform spectrum has a constrained peak, whereas the yellow platform was broader, spanning 500 - 600 nm. E) Normalized photoluminescence (PL) spectrum of the blue-fluorescent dye used in our clear coat paint under 405 nm laser excitation. The fluorescent dye PL spectrum closely matches that of the blue vane trap.

To process the 5.7 million camera trap images and maximize bumble bee detections, we compared the efficacy of two custom-built deep learning object detection models (Fig. 2A). One model used whole images at reduced resolution, whereas the tiling model used Slicing-Aided Hyper Inference (SAHI) to detect objects within overlapping subframes and aggregate predictions (Fig. 2 B, C). The tiling model exhibited better performance on a validation set of 296 images containing bumble bees, with higher recall and F1 scores across nearly all confidence thresholds (Fig. 2D). At a confidence level of 0.5, the tiling model achieved 96.8% recall and 98.1% F1, compared to 87.8% recall and 93.3% F1 for the whole-image model. The tiling model detected 57% more bumble bees (4,894 vs 2,776), without missing any found by the whole-image model. Furthermore, the tiling model outperformed a human reviewer, detecting 20% more images from a randomly chosen test day. However, the tiling approach was considerably slower, averaging 1.1 images per second compared with 88.5 images per second for the whole-image model.

**Figure 2:**
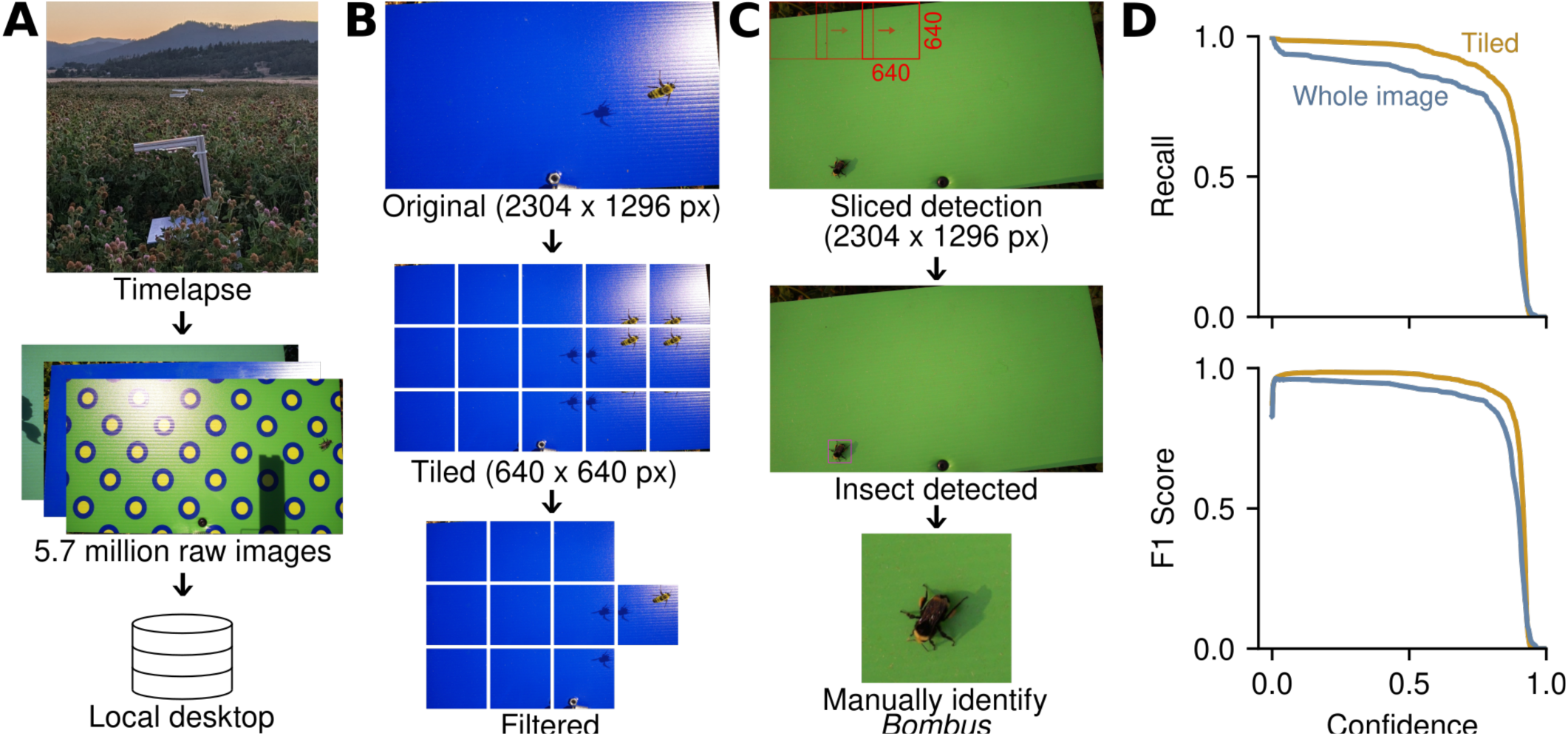
Tiling approach outperformed whole-image insect detection model. A) During the 18-day sampling period, images were captured at ∼1 Hz, resulting in about 5.7 million images. Images were transferred from the camera trap SD cards to local storage for processing. B) To train a tiling-based insect detection model, we split the original 2304 x 1296 px images into overlapping 640 x 640 px segments and removed partial insect fragments. C) We used Slicing-Aided Hyper Inference (SAHI) for insect detection, which processes overlapping 640 x 640 px frames across the entire image and then aggregates the predictions. From the detected insects we manually identified bumble bees. D) Recall-confidence and F1-confidence plots comparing the tiled (gold) and whole-image (blue-gray) models trained from YOLO11s weights. Recall measures the proportion of true positives correctly identified across confidence thresholds. The F1 score is the harmonic mean of recall and precision (the proportion of positive determinations that are correct), providing a balanced measure of model performance that penalizes both false positives and false negatives. The tiling model detected 57% more bumble bee images in the new camera trap data than the whole-image model.

### Bumble bee visits to distinct platform types

The 7,670 bumble bee images were recorded across all five platform types over 18 days. Daily detection counts (Fig 3A, top) varied independently of the amount of time cameras were operating each day (Fig 3A, bottom). Total camera runtime for each platform type ranged from 342.5 to 419.2 hours (blue and yellow platforms, respectively). To identify unique bee visits, we established a time-gap threshold by considering intervals between consecutive detections of the same *Bombus* species (c.f. Sittinger, 2024). We found that 93.8% of same-species, consecutive images occurred within 4 seconds, with none between 4 and 10 seconds (Fig. 3B). Therefore, we grouped same-species detections within 4 seconds, yielding 291 putative unique bumble bee visits. Across all traps, visitation frequency peaked between 9:00 and 10:00, though we recorded activity during all hours between 6:00 and 20:00 (Fig. 3C). We recorded the most visits to bullseye platforms, followed by green, fluorescent blue, blue, and yellow (95, 71, 54, 36, and 35 visits respectively; Fig. 3D).

**Figure 3:**
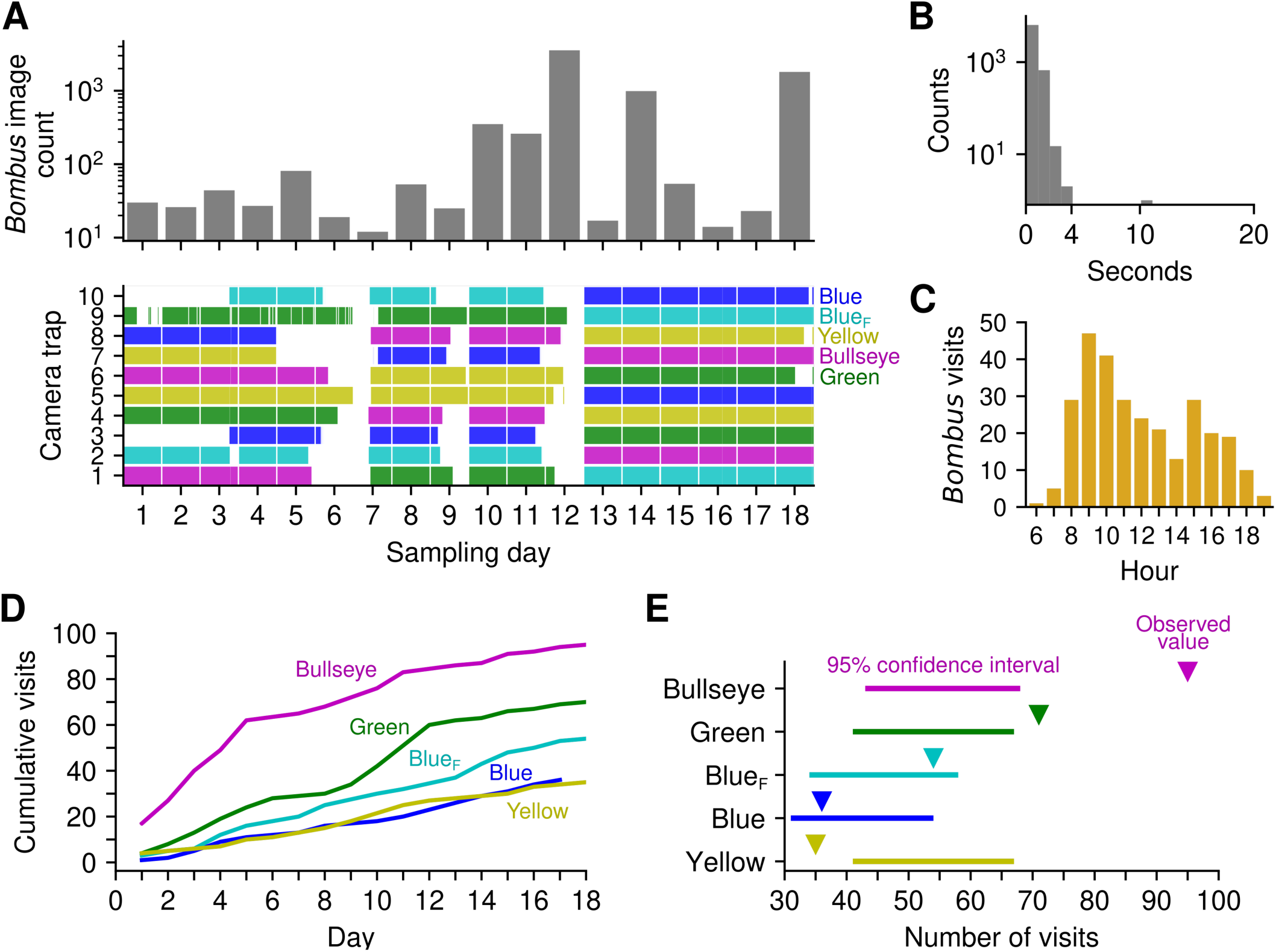
*Bombus* visitation across platform types. A) Daily counts of images containing bumble bees (top), with platform type and camera on-times (shaded segments) during each day’s sampling period (bottom). Across all panels, BlueF refers to platforms with blue ink with a fluorescent clear coat. B) Distribution of time intervals between consecutive images of the same bumble bee species. We used a 4 s threshold to define a unique visit; 93.8% of intervals were shorter than 4 s; none fell between 4 and 10 s. C) Distribution of bumble bee visits by time of day from 6:00 to 19:00. D) Cumulative bumble bee visit counts across platform types. E) Distributions of permutation test results from 10,000 iterations in which bumble bee visit times were randomly assigned to traps and counted if the corresponding camera was on at that time. Simulated visit counts were normally distributed for each platform type; horizontal lines indicate the 95% confidence interval and triangles mark observed visit counts.

We assessed whether the observed visit counts to distinct platforms differed from a null model of random visitation. To do this, we assigned all observed bumble bee visits to a random platform 10,000 times – counting visits only if the associated camera was recording. The observed visit counts to the bullseye and green platforms were significantly higher than this permuted distribution (bullseye, p < 0.001; green, p = 0.015; Fig. 3E). The fluorescent blue and blue platforms were not significantly different from the permuted distribution, and the yellow count was significantly lower (p = 0.004; Fig. 3E). After controlling for multiple comparisons using false discovery rate, visit counts to bullseye and green platforms remained significantly higher than chance (p < 0.001, p = 0.025, respectively), whereas yellow platforms stayed lower (p = 0.01). We used these randomly permuted data to test whether adding a fluorescent clear coat increased bumble bee visitation. The observed difference in visits between the fluorescent and blue platforms had a p-value of 0.058, suggesting any attractive effect of the fluorescent coat was modest. We observed no relationship between reapplication of fluorescent paint and visitation frequency. These findings suggest that the bullseye pattern was more attractive than any of its constituent colors.

Six identifiable *Bombus* species were detected: *B. vosnesenskii*, *B. fervidus*, *B. griseocollis*, *B. nevadensis*, *B. mixtus*, and *B. appositus* (Fig. 4A). Species composition and richness varied among platform types, with single visits of infrequent species driving differences (Fig. 4B). The two most common species, *B. vosnesenskii* and *B. fervidus*, were most frequently observed at all five platform types. Based on prior surveys, a small percentage (2-3%) of bees identified as *B. vosnesenskii* were likely *B. caliginosus*; the two species are difficult to distinguish using images or field observation (Rao and Stephen, 2009). Thirty bumble bee images were not identified to species. Green and yellow platforms were primarily visited by *B. vosnesenskii* and *B. fervidus*. *B. griseocollis*, the third most common species, was recorded on bullseye, blue, and fluorescent blue platforms. The highest species richness, of 5, was recorded by the bullseye and fluorescent blue platforms – which captured *B. nevadensis*, *B. mixtus* (bullseye platform only) and *B. appositus* (fluorescent platform only).

**Figure 4:**
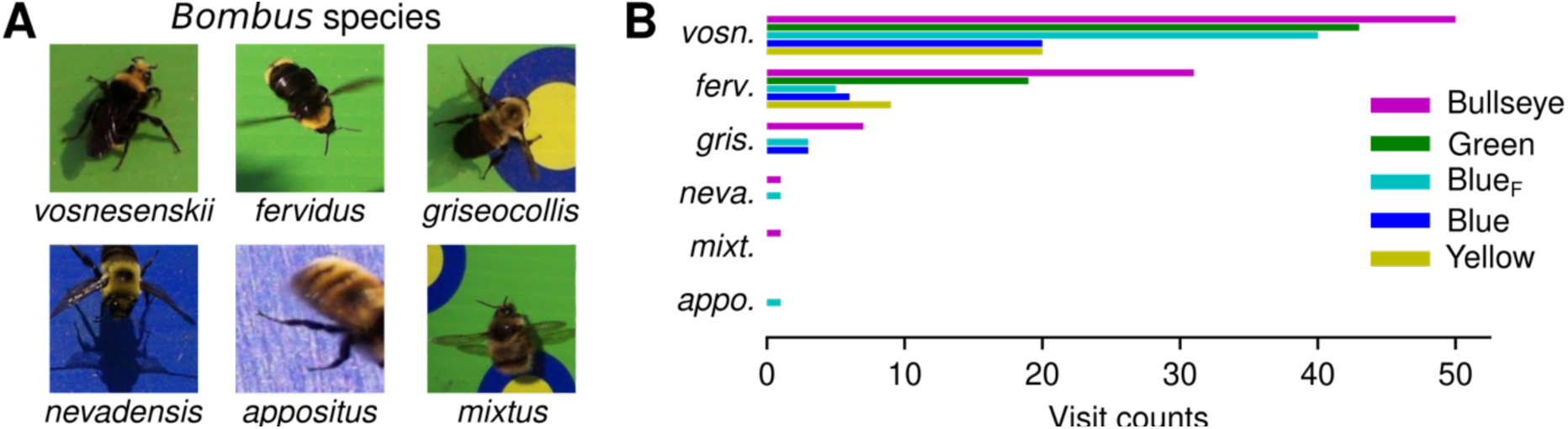
*Bombus* species visitation by platform. A) Cropped examples for each species of bumble bees detected on camera traps. B) Distribution of bumble bee species visits across platform types (Blue_F_ = blue with fluorescent clear coat).

### Comparison of distinct sampling methods

We evaluated capture rates from simultaneous sampling with blue vane traps, hand netting, and cameras (Fig. 5A). Over eight days, two blue vane traps collected 449 insects, including 130 bumble bees. Hand netting collected 176 bumble bees and 13 non-*Bombus* bees.

**Figure 5:**
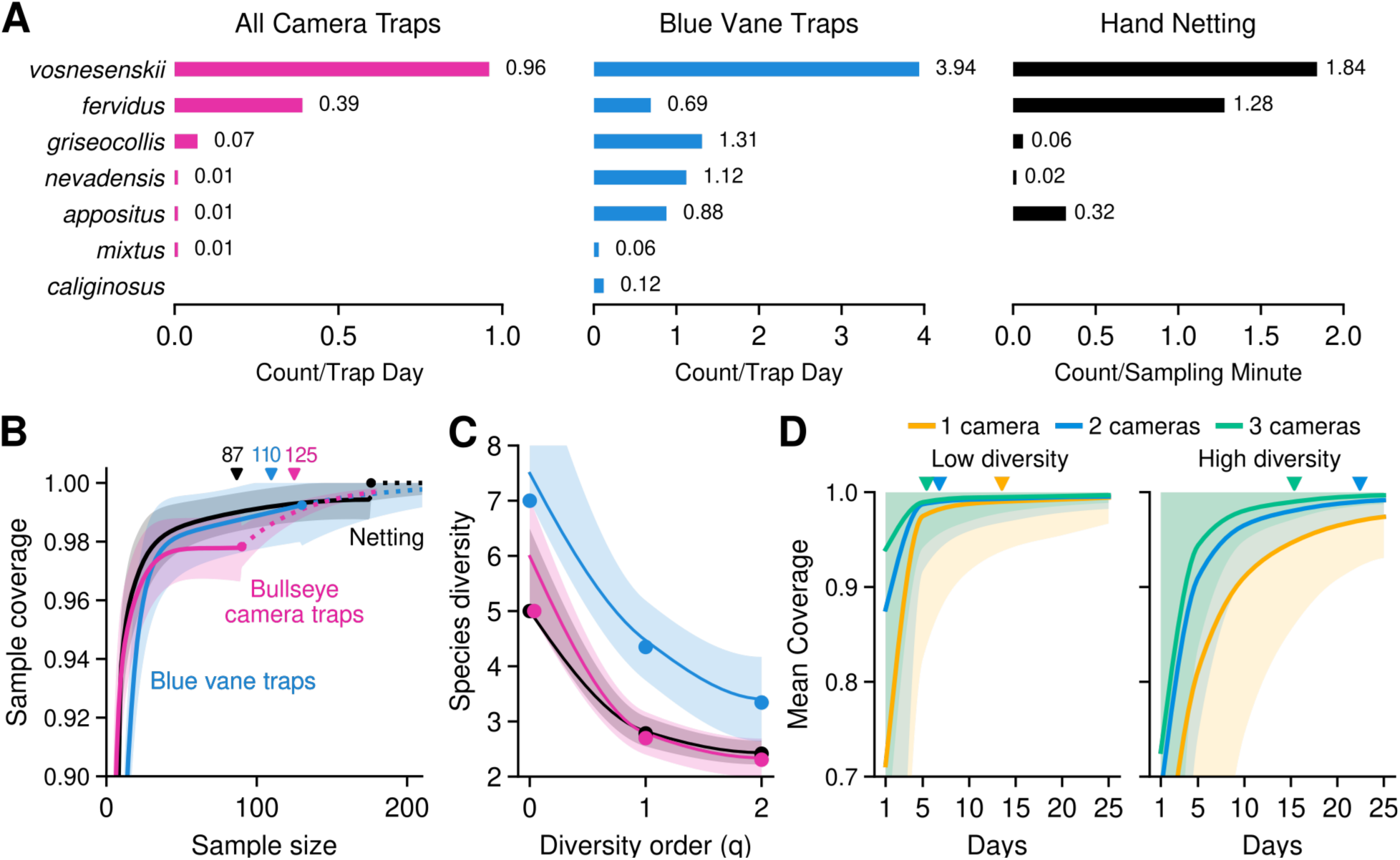
Comparison of capture rates, diversity estimates, and coverage across sampling methods. A) Capture rate per species for all ten camera traps, two blue vane traps, and 50 minutes of netting. Camera and blue vane trap rates were standardized as individuals per trap per day; netting was standardized as individuals per sampling minute. B) Sample coverage (the estimated proportion of individuals in the community represented by observed species) as a function of sample size for each method for blue vane traps (blue), netting (black), and two camera traps with bullseye platforms (pink). Solid lines show rarefaction; dotted lines show extrapolation; circle markers are observed values; shaded regions are 95% confidence intervals (CI). The small gap between the netting rarefaction and extrapolation curves reflects reduced coverage caused by downsampling in rarefaction. Triangles indicate the sample count at which each method reaches 0.99 coverage. Statistics were computed with *iNEXT*. C) Asymptotic diversity estimates across Hill numbers (q_0_ = richness, q_1_ = Shannon diversity, q_2_ = Simpson diversity) for each method; colors are the same as panel B. Asymptotic estimates predict the diversity expected with complete sampling. Circles mark observed values; solid lines show estimated values; shaded regions are 95% CI. D) Bootstrapped estimates of sample coverage over time for different numbers of camera traps at a low-diversity site (6 species; current study) and a high-diversity site (13 species; Oregon Bee Atlas data from Bend, OR). Rate of capture is modeled using the observed visitation rate to the bullseye platform. Shaded regions represent the 95% CI for each camera number; the upper bound of each CI is 1.0. Triangles mark when 0.99 coverage is reached.

Blue vane traps captured the highest bumble bee species richness, followed by camera traps and hand netting (7, 6 and 5 respectively). The most common bee, *B. vosnesenskii*, was collected 3.94 times per day on blue vane trap, compared to 0.96 times per day on camera traps. On average, the blue vane traps captured 7.8 bumble bees per trap each day, whereas hand netting yielded 3.5 bumble bees per person per ten-minute session. The camera traps together averaged 2.1 visits per sample day, whereas the bullseye platforms averaged 3.3 visits sample day.

We compared the efficiency of sampling methods by determining how coverage (the estimated proportion of individuals represented by observed species) varied with sample size (Fig. 5B). We used observed bumble bee data from the two bullseye traps to represent camera sampling. Netting was the most efficient at reaching 0.99 coverage, followed by blue vane traps and cameras (87, 110, and 125 samples respectively). We compared asymptotic diversity estimates across sampling methods to account for unequal effort. The blue vane traps outperformed the other methods across diversity orders, yielding the highest estimated richness, q_0_ = 7.5, evenness, q_1_ = 4.5, and diversity, q_2_ = 3.4 (Fig. 5C). Diversity estimates for netting (q_0_ = 5.0, q_1_ = 2.8, q_2_ = 2.4) and the two camera traps (q_0_ = 6.0, q_1_ = 2.8, q_2_ = 2.3) mostly overlapped, suggesting comparable performance. The higher richness estimate for the cameras suggests that additional species would likely be detected with increased sampling effort.

We simulated how camera-based sampling scales with effort using a bootstrapping approach to estimate coverage with varying numbers of camera traps and levels of bumble bee diversity. To model capture rates, we fit a negative binomial distribution to observed detection rates to estimate species counts across a 14-hour sampling day. We used the species abundance distribution from all camera traps in this study (6 species) to model a low-diversity site. We found that one camera reached 0.99 coverage after 13.5 days, whereas two and three cameras took 6.8 and 5.4 days respectively (Fig 5D, left). To model a high diversity site, we used data previously collected near Bend, OR, which included 13 species (Fig 5D, right). For this site, two cameras reached 0.99 coverage in 22.4 days, whereas three cameras took 15.4 days. One camera reached a maximum of 0.97 coverage after 25 days. These results suggest that deploying multiple cameras substantially accelerates reaching sampling completeness, especially at high-diversity sites.

## Discussion

In this study we developed and tested a custom, low-cost camera trap for continuous, non-lethal monitoring of diurnal insect pollinators (Figure 1). We trained deep-learning models to detect insects in images (Fig. 2A), and found that a tiling approach (Fig. 2B,C) improved bumble bee recall (Fig. 2D). We recorded bumble bee visits during all daylight hours (Fig. 3C). Bumble bee visitation was highest on platforms with a fluorescent bullseye pattern (Fig. 3E). In total, the camera traps recorded six bumble bee species, with the greatest richness on bullseye and fluorescent blue platforms. The overall capture rates of camera traps were lower than netting or blue vane traps (Fig. 5A); however, the camera trap estimates of bumble bee richness closely matched those of netting (Fig. 5B), while additionally providing time-stamped data unavailable in passive trap collections. Our simulation approach suggested adding additional cameras may yield substantial gains in sampling completeness at species-rich sites, with diminishing returns in low-diversity communities (Fig. 5D). These results demonstrate that camera traps can provide community-level assessments comparable to conventional lethal sampling, offering a scalable, non-invasive route toward long-term monitoring of vulnerable bee populations.

Platforms with a repeating bullseye pattern, which mimicked a floral contrast cue, attracted more bumble bees than uniform color platforms. This finding (Fig. 3E) is consistent with prior work showing that naturally occurring bullseye flower patterns are attractive to bees (Horth et al., 2014; Koski and Ashman, 2014), and that highly contrasting colors can attract bees on two-dimensional platforms (De Jager et al., 2017; Lunau et al., 2025). Prior studies have used concentric rings on platforms with captive bees (Hunt and Chittka, 2015; Rohde et al., 2013); here we show that bullseye patterns, mimicking floral cues, are attractive to wild bumble bees. Of the constituent colors that comprise the bullseye pattern; the solid green platform was the most visited (Fig. 3E). This high attractiveness of a uniform green lure differs from prior studies (Acharya et al., 2021; Ostroverkhova et al., 2018; Sipolski et al., 2019). Whereas color contrast is typically a short-range stimulus, green contrast cues have been shown to play a role in long-distance spatial processing which may have contributed to this unexpected response (Chittka and Tautz, 2003; Giurfa et al., 1996). The large green background and high-contrast concentric circles of the bullseye platforms potentially integrate both long-range and short-range cues; together, they may have elicited bumble bee approaches. The modest effect of adding a blue-fluorescent coat (Fig. 3E) contrasts with the high attractiveness of blue vane traps, which may benefit from strong color specificity and the vertical vane orientation (Hall, 2018; Stephen and Rao, 2005). Compared to the blue vane traps, the lower spectral purity of our blue ink (Fig. 1D) may have weakened the attractive signal (Lunau and Dyer, 2024). The low visitation to yellow platforms is consistent with prior studies (Sircom et al., 2018; Stephen and Rao, 2007). The high bumble bee recruitment to bullseye platforms and blue vane traps points to contrast patterns and spectrally pure fluorescent colors as promising drivers of trap attractiveness. The relative attractiveness of the bullseye pattern’s distinct features – such as color combinations, fluorescence, pattern arrangement, and spatial orientation – could be studied further with these camera traps.

By imaging continuously, we recorded bumble bee visits throughout all daylight hours (Fig. 3C). This suggests that full characterization of bumble bee communities, with sensitivity to rare species, may require full-day monitoring; this should be accounted for when designing camera trap power systems. The runtime of our traps was constrained by finite battery life, which could be readily addressed by adding solar power. The principal limitation of our time-lapse imaging approach is the steady accumulation of non-target images, which limits storage capacity. For long-term, automated monitoring, it may be critical to capture images selectively when insects are present. Recent studies have triggered image capture using AI-enabled cameras running lightweight detections model in real-time (Gardiner et al., 2025; Sittinger et al., 2024). This approach produced fewer images, thereby reducing downstream processing. We found that a tiled inference model improved bumble bee detection – identifying 57% more images than the whole-image approach (Fig. 2D) – but was 80 times slower. Going forward, combining coarse, on-device initial filtering of insects followed by precise, tiling-based localization offline could be key to balancing accuracy with practical, long-term operation.

We found that each sampling method produced distinct estimates of the local bumble bee community. Blue vane traps collected large numbers of bumble bees, with the highest estimated species richness and diversity, consistent with prior studies (Fig. 5B; Hall, 2018; Kimoto et al., 2012; Stephen and Rao, 2005). Hand netting achieved full sample coverage with the fewest specimens but yielded the lowest species richness, consistent with studies suggesting that apparent completeness in sampling may fail to detect rare species (Fig. 5B, C; Benoit et al., 2018; Fisher et al., 2022; Sgarbi et al., 2020). Although active netting allows targeted, rapid accumulation of specimens, it is constrained by observer skill, weather, and limited temporal coverage, often resulting in biased detection toward abundant or easily captured species (O’Connor et al., 2019; Popic et al., 2013). Our two most attractive traps, with bullseye platforms, matched netting in observed species richness despite capturing fewer specimens.

These two traps also yielded higher estimated richness than netting (6 vs 5). The blue vane trap also yielded an estimated richness greater than observed (7.5 vs 7), which is relevant considering that we did not detect *B. californicus* but it has been observed previously at the same site (Rao and Stephen, 2009). Although the camera traps required more samples to reach comparable coverage to other methods, our simulations show that increasing unit number substantially improves sampling efficiency, particularly at high-diversity sites (Fig. 5D). The scalability of camera traps may therefore be key to reaching recruitment levels necessary for detecting rare species, which can require extensive sampling in diverse areas (Fisher et al., 2022).

After image collection, taxonomic identification remains a major bottleneck, requiring time and expertise. Deep-learning classifiers can achieve species-level identification (Sittinger et al., 2024; Spiesman et al., 2021), though model generalization across camera trap systems can be limited. Cryptic species (e.g. *B. vosnesenskii* and *B. caliginosus*) further challenge image classification; additional data inputs such as acoustics (Gradišek et al., 2017; Kohlberg et al., 2024) or eDNA (De Jager et al., 2017; Jones et al., 2025) may be necessary to improve taxonomic fidelity. Expanding image annotation beyond *Bombus* to include other bee genera and pollinator guilds would also enable richer comparisons among detection methods. Continued refinement of on-board detection and post-processing classification will make camera traps increasingly capable for long-term remote monitoring, or other uses, such as mark-and-recapture navigation experiments.

Our study contributes to a growing body of work exploring the potential for camera traps as an insect survey method (Besson et al., 2022; Bjerge et al., 2023; Droissart et al., 2021; Simokat et al., 2024; Sittinger et al., 2024). Unlike netting, camera traps operate continuously, are non-invasive, and can be deployed at scale; unlike vane traps, image-based sampling is nonlethal and provides precise visit times that can be leveraged for analysis of species-specific activity patterns. Distributing a network of camera traps across a landscape could fill gaps in the data patchwork of species presence, climate niche occupancy, and community associations, providing a more comprehensive view of pollinator health and ecology. Continuous, spatially extensive monitoring of bees would improve detection of rare and endangered species and help link insect activity with environmental variables. With continued refinement of visual lures and detection algorithms, camera traps could offer a robust, non-invasive alternative to traditional lethal methods.

## Acknowledgments

We thank M. Gragg and J. Culley for collecting spectral data from the platforms, and taxonomists August Jackson and Brooke Ruby for identifying specimens and insects in images.

## Notes

### Competing Interest Statement

The authors have declared no competing interest.

### Funding

- USDA/NIFA AFRI: 05857
- USDA-ARS: 59-2072-3-025

